# Liquid and vapor-phase antifungal activities of natural borneol against Candida albicans and its germ tube formation

**DOI:** 10.1101/2021.03.02.433682

**Authors:** Yazhou Wang, Huiling Liu

**Affiliations:** changzhou cancer hospital

**Author notes:** Correspondence author: yazhou wang.

**Keywords:** Antifungal effects, Candida albicans, Germ tube, Natural borneol

## Abstract

Candida albicans infection mainly occurs in patients with suppressed immune function, and it is also the main pathogen of hospital infection. The new strategies are needed to treat the existing resistance of antifungal drugs. The use of natural products aimed at controlling fungal diseases is considered an interesting alternative to synthetic fungicides due to their lower adverse reactions, lower cost to plant preparations compared to modern conventional pharmaceuticals. Natural borneol has a long history of treating ulcers and local infections. In this study, the minimum inhibitory concentration of natural borneol on ATCC10231 and 10 clinically isolated Candida albicans was determined by vapor phase method and dilution method, and the influence of sub-minimum inhibitory concentration on the formation of Candida albicans hyphae was observed. We found that the minimum inhibitory concentration of ATCC10231 and 10 clinically the isolates in the vapor phase were both 0.4 mg/cm^3^, agar and broth dilution methods were 2 mg/mL. The vapor phase of natural borneol has a better inhibitory effect on Candida albicans, Sub-mic concentration of borneol (0.125-1mg/ml) in the liquid phase inhibits the 60%-99% formation of Candida albicans germ tube. Natural borneol is a potential natural medicine for the treatment and prevention of Candida albicans infection. It brings new insights into the development of novel effective antifungal drugs.

## 1. Introduction

Candida albicans usually colonize the gastrointestinal, reproductive tract, oral cavity, and skin (1,2). It exists in single-celled sprouting yeast, pseudomycelia, and filamentous forms, and can transform between the three forms (3,4). The use of antibiotics, microbial infections, immunosuppressive therapy, anti-cancer treatment, AIDS, diabetes, etc., which reduce the host immunity and cause the overgrowth of Candida albicans to cause mucosal infection or systemic infection (5). Candidiasis and disseminated candidiasis have a mortality rate of 40%(6) and are the second leading cause of death from nosocomial infections(7,8). Antifungal drugs used to treat Candida albicans have developed drug resistance (9). Therefore, new treatment strategies are needed to combat the infection of Candida albicans.

The use of natural products aimed at the control of fungal diseases is considered an interesting alternative to synthetic fungicides due to their lower adverse reactions, lower cost to plant preparations compared to modern conventional pharmaceuticals(10). Many essential oils have been demonstrated to exert biological activity in vivo and in vitro, which has justified research on the characterization of their antimicrobial activity. Natural borneol (NB) is one of the components of essential oil which has antifungal activity. NB also is a time – honored herb in traditional Chinese medicine. It is an almost pure chemical with a chemical composition of (+) – borneol. which has a long history in clinical applications to treat burns, sprains, pain, ulcers. Studies have found that borneol promotes wound healing(11,12) and has a good local analgesic effect (13), as a skin penetration enhancer, promote drug penetration into the skin (14). This study examines whether NB in the liquid phase and vapor phase can inhibit the Candida albicans. We also investigated the effect of NB on the formation of Candida albicans germ tube.

## 2. Materials and Methods

### 2.1 Chemicals and strains

Natural borneol was purchased from the Xinhuang Natural borneol, Hunan, China. Inc. Candida albican ATCC10231 was provided by Anto company, Henan, China. 10 clinical isolates were obtained from the clinical laboratory of Changzhou cancer hospital, Changzhou, China. Sabouraud dextrose broth (SDB) and Sabouraud dextrose agar(SDA) were purchased from Hangzhou Tianhe. Hangzhou, China. The strains were stored in 20% glycerol broth at - 80°C refrigerators. Before using all strains were cultured on SDA at 35°C for 24 hours and then suspended in 2 mL of 0.85 % normal saline to 0.5 McFarland standard.

### 2.2 **Determination of borneol anti-Candida** albicans **activity by Vapor Phase test**

The effect of Natural Borneol Vapor Phase on candida albicans was studied with invert Petri dishes method as previously reported with some modifications(15).The SDA was poured into a 60 mm Petri dish (PD) and 100 μl of the strain suspension was evenly spread on the SDA. The borneol concentration of 5.4, 10.8, 21.6mg/ml was prepared with anhydrous ethanol. 500 μL of each borneol solution was distributed on the surface of a sterile filter paper disc with a diameter of 60 mm. The disc was placed on the plate cover after solvent evaporation. PDs were hermetically closed with Parafilm adhesive tape to avoid evaporation, and then were incubated at 37 °C for 24 h. After incubation, filter papers were removed and screen the minimum inhibitory concentration (MIC). The MIC was determined by comparison with the control and was defined as the lowest concentration of borneol inhibiting the visible growth, expressed by mass/plate space volume, i.e. mg/cm^3^. The space inside of the sealed Petri dish was calculated to be 27 cm^3^ air. The filter paper treated with absolute ethanol without borneol as the control. All tests were repeated at least three times.

### 2.3 Determine the MIC of borneol by micro broth dilution method

The effect of Natural Borneol was studied with Micro broth dilution method as previously reported with some modifications(16). The 0.5 McFarland standard stains were 1: 1000 dilution with Sabouraud dextrose broth, then take 100μl into 96-well plates. 8 mg/mL borneol were made with Sabouraud dextrose broth, added Tween 80 (5%, v/v) to enhance the borneol solubility. After this, a serial twofold dilution was prepared from 4, 2 mg/mL. 8, 4, 2 mg/mL borneol solution added 100 μl to 96-well plate. 37 °C culture for 24 hours. The final concentration of borneol was 4, 2, 1 mg/mL. SDB media containing 5% of Polysorbate 80 were used separately as controls. and all tests were repeated three times. The MIC value is the concentration of borneol without visible growth.

### 2.4 Determine MIC of borneol by agar dilution method

MICs of borneol were determined by agar dilution assay(16). The agar plates were prepared by adding SD containing various concentrations of borneol (i.e. 1, 2, 4 mg/mL). Tween 80 (5%, v/v) was added to enhance the borneol solubility. plates were inoculated with 10^3^ cfu using the inoculate of Candida strains prepared as above. Plates were kept in triplicates for each concentration. Plates with Tween 80 but without any borneol were used as control. All these plates were incubated at 35 °C. The MIC values were determined as the lowest concentration of borneol preventing visible growth of Candida strains.

### 2.5 Screening effects of Natural Borneol Sub-MIC on Candida Albicans germ tube formation

The assay was then carried out as previously described by ALVES M ea al.(16), the density of the Candida albicans suspension was adjusted to 10^6^ CFU/mL, and 990 μL of the strains solution was distributed to test tubes, and 10 μL of borneol of different concentrations was added to each test tube and incubated at 37°C for 3 hours. The final borneol concentration is 1, 0.5, 0.25, 0.125, 0.063, 0.031 mg/mL. Then, record the formation of sprout tubes under an optical microscope (40×). 1 % (v/v) Tween 80 was used as the control. The result is the mean ±standard deviation (SD) of three independent experiments

### 2.6 Statistical Analysis

Data are expressed as mean ± standard error (SEM) of the mean. Statistical significance was determined using one-way analysis of variance (ANOVA), followed by Dunnett’s post hoc test. The statistical analysis was performed using SPSS19. Differences were considered significant for p < 0.05.

## 3. Results

### 3.1 Screening of borneol for anti-Candida albicans activity

The MIC of borneol is presented in table 1. Overall, vapor phase borneol (MIC 0.4 mg/cm^3^) inhibited candida albicans more effectively than in liquid(MIC 2 mg/mL).

**Table 1.** Antifungal activity of borneol against Candida albicans.

| Strain | vapor phase<br>test MIC<br>(mg/cm<sup>3</sup>) | Agar dilution<br>method MIC<br>(mg/mL) | Micro Broth dilution<br>method MIC<br>(mg/mL) |
| --- | --- | --- | --- |
| ATCC10231 | 0.4 | 2 | 2 |
| 995020 | 0.4 | 2 | 2 |
| 0556696 | 0.4 | 2 | 2 |
| 056162 | 0.4 | 2 | 2 |
| 9452193 | 0.4 | 2 | 2 |
| 0466435 | 0.4 | 2 | 2 |
| 239054 | 0.4 | 2 | 2 |
| 238089 | 0.4 | 2 | 2 |
| 238094 | 0.4 | 2 | 2 |
| 238449 | 0.4 | 2 | 2 |
| 238350 | 0.4 | 2 | 2 |

### 3.2 Effects of Natural Borneol Sub-MIC Concentration on Candida Albicans germ tube formation

As shown in fig 1, borneol inhibited the germ tube formation at concentrations below its MIC, Candida albicans germ tube decreased by 99%, 97%, 78%, 60% in 1, 0.5,0.25,0.125mg/ml NB respectively.

**Figure 1.**
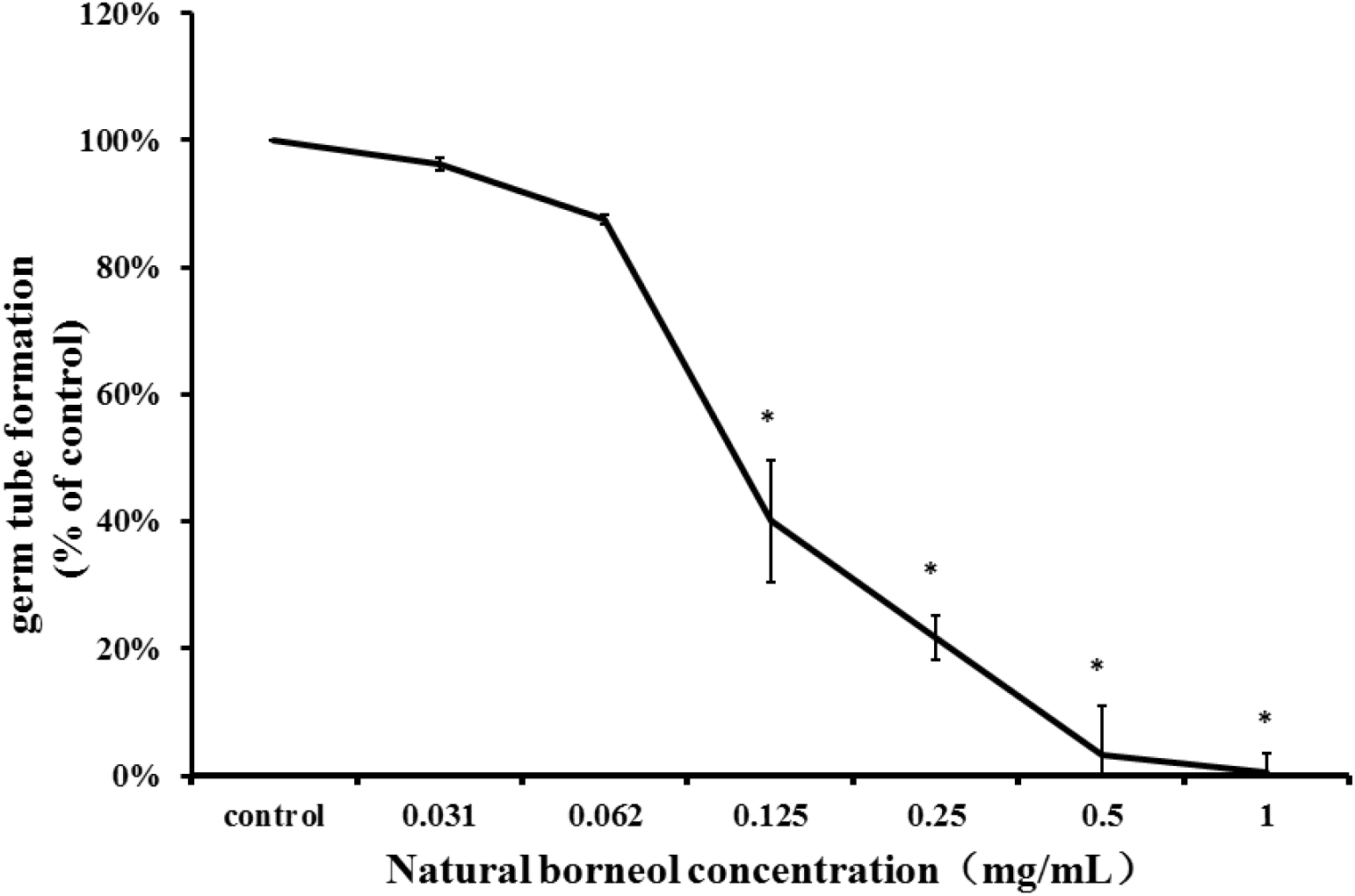
The effect of sub-MIC of natural borneol on Candida albicans germ tube. * *P* <0.05

## 4. Discussion

It is known that the antimicrobial action of essential oil depends on their presence in gaseous form facilitating their solubilization in cell membranes(17).Amit K Tyagi et al. observed that minimum inhibitory dose of lemon grass essential oil was significantly lower than peppermint essential oil and eucalyptus essential oil in gaseous phase, leading to deleterious morphological changes in cellular structures and cell surface alterations(18). We found that the anti-Candida albicans activity of natrual borneol MIC in vapor phase (0.4 mg/cm^3^) was significantly lower than in liquid phase (2mg/mL). The anticandida activity may depends upon the diffusability and solubility of the borneol into the agar in liquid phase while the activity of the vapour assay depends upon the volatility of borneol. Mycelium is the critical factor for the virulence of Candida albicans and biofilm formation. The mycelia form of Candida albicans is more invasive than the yeast form (19). Candida albicans invade the mucosa cause mucosal infection or breaks through the mucosa cause Candidiasis and disseminated candidiasis.

The mycelia kill the phagocytes and allow the yeast cells to escape (20). Therefore, the development of antifungal drugs for the biological process of mycelia transformation is one way to overcome drug resistance, preventing it change from a harmless symbiosis state to a pathogen, maintaining the symbiosis state rather than eradicating it is a potential strategy (21). In this study, the effect of the borneol on the germ tube formation in C. albicans ATCC 10231 was assessed. Sub-mic concentration of borneol (0.125-1mg/ml) in liquid phase inhibit the 60%-99% formation of Candida albicans germ tube.

Despite the differences between the vapor and solution methods, our results demonstrate that NB is very active on Candida albicans. It brings new insights on the development of novel effective antifungal drugs.

However, further studies are required to study the effect and toxicity of NB in in vivo and to establish if they could be safely used as antifungal agents.

